# Effects of panitumumab on the proteome of treated EGFRvIII glioblastoma cells and on their extracellular vesicles

**DOI:** 10.64898/2026.10.22.760940

**Authors:** Amy Gateau, Baptiste Moeglen-Paget, Stéphanie Durand, Hussein Akil, Georges Humbert, Barbara Bessette, Fabrice Lalloué

**Affiliations:** UMR INSERM 1308 CAPTuR, Faculty of Medicine, University of Limoges, Limoges, France; XLIM Research Institute, UMR 7252 CNRS/University of Limoges, 123 Avenue Albert Thomas, 87060 Limoges, France

## Abstract

Glioblastoma (GB) is an aggressive brain tumor for which reliable biomarkers of therapeutic response are still needed. Extracellular vesicles (EVs) can carry molecular components derived from their parent cells, thereby providing information on changes associated with the tumor and treatments. Among potential biomarkers, epidermal growth factor receptor variant III (EGFRvIII), is found in more than 40% of GB patients and is associated with aggressive tumor behavior and therapeutic resistance. EGFRvIII has also been detected on the surface of EVs, suggesting its potential value for monitoring tumor-associated molecular changes. This study investigated the effects of panitumumab, an anti-EGFR monoclonal antibody, on the proteomic profiles of EVs derived from EGFRvIII cells. U87 and U87-EGFRvIII cells and their derived EVs were characterized before and after treatment. EGFRvIII expression in cells and EVs was assessed by Western blotting and flow cytometry. Proteomic analyses were performed on cell lysates and EVs before and after treatment. Our results demonstrate that panitumumab significantly reduced surface EGFRvIII on EVs derived from U87-EGFRvIII cells, whereas no significant decrease was observed in cell lysates. This suggests that treatment may affect EGFRvIII trafficking or its incorporation into EVs. The proteomic analysis highlighted much more changes in EVs than in their parental cells after treatment, particularly in proteins associated with extracellular matrix organization and collagen. Together, these results demonstrate that panitumumab treatment is associated with changes in protein profile from EVs derived from EGFRvIII cells suggesting that the panitumumab treatment might have an impact on cells that could be reflected on EVs released by glioblastoma cells. In conclusion, our results suggest that EVs derived from EGFRvIII cells could serve as a useful circulating biomarker for assessing treatment efficacy and for preventing treatment escape.

## 1. Introduction

The epidermal growth factor receptor variant III (EGFRvIII) is a tumor-associated alteration found in around 40% of glioblastoma (GB), resulting from an in-frame deletion of exons 2-7 of the *EGFR* gene. This truncated receptor has a constitutive signaling activity and contributes to oncogenic processes, notably cell proliferation, survival, angiogenesis and invasion ^1–3^. Beyond its expression in tumor cells, EGFRvIII has been detected in extracellular vesicles (EVs) released by glioma cells, raising interest in EV-associated EGFRvIII as a potential biomarker for liquid biopsy ^3,4^. Indeed, EVs carry proteins and other molecular components derived from their cells of origin and can be isolated from biological fluids, making them attractive candidates for investigating tumor-associated molecular changes. However, the presence of EGFRvIII in EVs and its potential as a marker of treatment-associated changes require further investigations.

Targeting EGFR is a therapeutic strategy commonly used in several cancers, even though its efficacy depends on the tumor type and molecular context. Panitumumab is a fully human monoclonal antibody directed against the extracellular domain of EGFR, currently used in patients with RAS wild-type metastatic colorectal cancer ^5^. Its effects on EGFR-related proteins in GB, in particular concerning EGFRvIII in EVs, remain insufficiently characterized. In particular, it is important to determine whether the changes induced by the treatment in cells expressing EGFR can be reflected in the EVs they release and if these changes affect the detection of EGFRvIII at the EVs surface. By investigating these questions, we could clarify the relationship between cellular treatment responses and the molecular characteristics of tumor-derived EVs.

In addition to proteins specifically associated with the tumor, the total protein composition of the EVs could also provide information about cellular changes associate with the treatment. Therefore, a comparison of the proteomic profiles of cells and their derived EVs before and after treatment could help to identify biological pathways associated with the cellular response of panitumumab and also determine if these changes are reflected in the EV proteome.

In this study, we investigated the effect of panitumumab on two glioblastoma cell lines, specifically the U87 cells, which served as our control, and the U87-EGFRvIII cells, which overexpress EGFRvIII, as well as on the EVs extracted from these cells. We aimed to evaluate potential changes in the abundance of EGFR or EGFRvIII, the presence of EGFRvIII on the surface of the vesicles, or changes in the characterization of the vesicles following treatment. We therefore performed a comparative proteomic analysis of cellular and vesicular lysates before and after treatment to identify changes in the abundance of certain proteins. Combining targeted EGFRvIII analyses with global proteomic profiling allows us to study the influence of panitumumab on the molecular characteristics of glioblastoma-derived EVs and to identify potential signaling pathways involved in these changes.

## 2. Materials and methods

### 2.1 Cell culture

The human glioblastoma cell line U87-MG was cultured in DMEM GlutaMAX medium (Gibco) supplemented with 10% heat-inactivated fetal bovine serum (FBS; 56°C for 30 min), 1% penicillin–streptomycin (Gibco), 1% sodium bicarbonate (Gibco), and 1% non-essential amino acids (Gibco). The human glioblastoma cell line U87-EGFRvIII, which overexpresses EGFRvIII, was cultured in MEM medium (Gibco) supplemented with 10% FBS, 0.2% gentamicin (#15710064, Thermo Fisher Scientific), 200 µg/mL G418 (ant-gn-2, InvivoGen), and 1% penicillin–streptomycin. Cells were maintained at 37°C in a humidified atmosphere containing 5% CO₂.

### 2.2. Extracellular vesicle obtained from cell lines

U87-MG and U87-EGFRvIII cells were seeded at a density of 1.7 × 10⁶ cells in T175 flasks containing 18 mL of culture medium. The following day, the culture medium was replaced with EV-depleted medium. This medium was prepared as described above, except for the FBS used. FBS was depleted of nanoparticles by ultracentrifugation at 120,000 × g for 17 h at 4°C, followed by filtration through a 0.2 µm filter. The cells were then cultured for 72 h, after which the conditioned media were collected.

### 2.3. Extracellular vesicle isolation by ultracentrifugation

The conditioned media were collected and subjected to an initial centrifugation at 300 × g for 10 min to remove dead cells, followed by centrifugation at 2,000 × g for 10 min to remove cellular debris. The resulting supernatants were then filtered through a 0.2 µm filter to preferentially remove larger particles, including large microvesicles. The samples were subsequently ultracentrifuged using an Optima XPN-80 ultracentrifuge (Beckman Coulter) at 120,000 × g for 2 h. This ultracentrifugation step was performed twice to pellet and wash the extracellular vesicles.

### 2.4. Extracellular vesicle characterization by nanoparticle tracking analysis (NTA)

NTA measurements were performed using a NanoSight NS300 instrument (Malvern Panalytical Ltd.). Data acquisition and analysis were performed using the integrated NanoSight NTA software (version 3.3.301, Malvern Panalytical Ltd.). The camera level was set to 14, and the detection threshold was set to 5. Samples were diluted in PBS to a final volume of 500 µL, with the concentration adjusted to obtain approximately 50 particles per frame (30–80 particles per frame). For each measurement, five 60-s videos were recorded at a cell temperature of 25°C and a syringe pump speed of 22 µL/s (100 arbitrary units). EVs were detected using a 488-nm blue laser and a scientific CMOS camera. The mean particle size, mode size, and particle concentration (particles/mL) were determined using the NTA software.

### 2.5. Protein extraction

Cells were detached using trypsin and centrifuged at 200 × g for 7 min. Cell pellets were then washed with PBS (Gibco) and lysed using 100 µL of 1× Cell Lysis Buffer (#9803, Cell Signaling Technology) per 1 × 10⁶ cells. The lysis buffer was supplemented with 1% protease inhibitor cocktail and 1% phosphatase inhibitor cocktail (#P0044-5ML, Sigma-Aldrich). Chemical lysis was performed on ice for 30 min. Cell lysates were subsequently sonicated using a Vibra-Cell Sonifier and centrifuged at 18,000 × g for 20 min at 4°C. The resulting lysates were then subjected to protein quantification using the Bradford assay.

### 2.6. Western blot

Proteins were denatured at 95°C for 5 min in 4× Laemmli buffer containing 10% β-mercaptoethanol, separated on 12% polyacrylamide gels, and transferred onto membranes using the Trans-Blot Turbo Transfer System (Bio-Rad). Membranes were blocked with 5% BSA in TBS-T for 1 h at room temperature and incubated overnight at 4°C with the appropriate primary antibodies. After washing with TBS-T, membranes were incubated with HRP-conjugated secondary antibodies (Dako, Agilent; 1:1000). Protein detection was performed using Immobilon ECL Ultra Western HRP Substrate (Merck) and an Azure imaging system (Azure Biosystems). TotalStain Q (Azure Biosystems) was used for protein normalization and quantification.

| Target | Species | Reference | Supplier |
| --- | --- | --- | --- |
| CD81 | IgG Mouse | sc-166029 | Santa Cruz |
| CD9 | IgG Rabbit | ab307085 | Abcam |
| TSG101 | IgG Rabbit | ab125011 | Abcam |
| Calnexin | IgG Rabbit | 2433S | Cell Signaling |
| EGFR | IgG Rabbit | 4267S | Cell Signaling |
| p-EGFR | IgG Rabbit | 3777S | Cell Signaling |
| EGFRvIII | IgG Rabbit | MA5-36216 | Invitrogen |
| HSP90 | IgG Rabbit | 4877S | Cell Signaling |

### 2.7. Panitumumab treatment

According to the experimental conditions, cells were treated with 10 µg/mL panitumumab (Vectibix, Amgen, Thousand Oaks, CA, USA), provided by Limoges University Hospital (CHU de Limoges). To assess the effect of panitumumab on the EGFR/EGFRvIII axis, cells were seeded at a density of 1.7 × 10⁶ cells in T175 flasks. After treatment, conditioned media were collected 72 h later, and extracellular vesicles were isolated according to the protocol described above.

### 2.8. Flow cytometry analysis of extracellular vesicles

#### 2.8.1. Magnetic beads functionalization

For flow cytometry analysis, commercially available magnetic beads presenting CD63 (Exosome-Human CD63 Isolation/Detection Reagent, #10606D, Invitrogen) were used as a positive control. Dynabeads Protein A (#10002D, Invitrogen) were resuspended by vortexing for 30 s. Subsequently, 50 µL of beads were transferred to a 2-mL tube and placed on a magnetic rack to remove the supernatant. The beads were then resuspended in 200 µL of PBS containing 0.02% Tween-20, supplemented with 1 µg of the antibody of interest, and incubated under rotation for 10 min at room temperature. The supernatant was subsequently removed using the magnetic rack, and the beads were resuspended in 200 µL of PBS containing 0.02% Tween-20.

| Target | Species | Reference | Supplier |
| --- | --- | --- | --- |
| CD63 | IgG Rabbit | Ab68418 | Abcam |
| EGFRvIII | IgG Rabbit | MA5-36216 | Invitrogen |
| HSP90 | IgG Rabbit | 4877S | Cell Signaling |
| Isotype Rabbit | IgG Rabbit | 14-4616-82 | Invitrogen |

#### 2.8.2. PKH67 labeling of extracellular vesicles

PKH67 from a commercial kit (Green Fluorescent Cell Linker Kit; Sigma-Aldrich) was used to label EVs produced by U87-MG and U87-EGFRvIII cells. Following the first ultracentrifugation at 120,000 × g, the EV pellet was resuspended in 250 µL of Diluent C. In parallel, 1 µL of PKH67 was diluted in 250 µL of the same diluent, and this solution was subsequently added to the EV suspension. After 5 min of incubation at room temperature, the labeling reaction was stopped by adding 1 mL of EV-free culture medium for 2 min. To wash the sample, 7 mL of 1× PBS were added, followed by a second ultracentrifugation at 120,000 × g for 2 h at 4°C. Finally, the EV pellet was resuspended in 100 µL of 0.1% BSA in PBS and stored at 4°C.

#### 2.8.3. Immunocapture of extracellular vesicles using magnetic beads

A 10-µL aliquot of functionalized magnetic beads was washed with 0.1% BSA in PBS using a magnetic rack. After removal of the supernatant, 50 µL of PKH67-labeled EVs were incubated with the beads in the presence of an additional 50 µL of 0.1% BSA in PBS, with shaking overnight at 4°C. The following day, the beads were washed twice with 0.1% BSA in PBS and resuspended in 200 µL of PBS. The fluorescence signal was then detected using a CytoFLEX flow cytometer (Beckman Coulter). Data were analyzed using Kaluza software (Beckman Coulter).

### 2.9. Proteomic analysis

#### 2.9.1. Sample preparation

Total protein lysates were extracted from U87-MG and U87-EGFRvIII cell lines, as well as from extracellular vesicle samples secreted by these cell lines, as previously described in 2.5 section using a commercial lysis buffer from Cell Signaling Technology (20 mM Tris-HCl pH 7.5, 150 mM NaCl, 1 mM Na₂EDTA, 1 mM EGTA, 1% Triton, 2.5 mM sodium pyrophosphate, 1 mM β-glycerophosphate, 1 mM Na₃VO₄, 1 µg/mL leupeptin, and 1 mM PMSF). All lysates were incubated on ice for 30 min. Cell lysates were then briefly sonicated and centrifuged at 18,000 × g for 20 min.

Prior to digestion, total protein concentrations in EV samples were determined using the Pierce BCA Protein Assay Kit (Thermo Fisher Scientific), while cellular protein concentrations were determined using the Bradford assay. An equal amount of total protein was used for subsequent proteomic analysis for all samples. This normalization step ensured that any differences observed in the abundance of detected proteins reflected variations in EV composition rather than variability in EV recovery.

Proteomic analyses were performed at the Proteomics Platform of the Institute of Genetics and Molecular and Cellular Biology (IGBMC), 1 rue Laurent Fries, 67404 Illkirch Cedex, France.

#### 2.9.2. Liquid digestion

Protein mixtures were TCA-precipitated overnight at 4°C, pellets were washed twice with 1 mL cold acetone, dried and dissolved in 2 M urea in 0.1 mM Tris-HCl pH 8.5 for reduction (5 mM TCEP, 30 min.), and alkylation (10 mM iodoacetamide, 30 min.). Trypsin digestion was carried out at 37°C and overnight. Peptide mixtures were then desalted on C18 spin-column and dried on Speed-Vacuum before LC-MS/MS analysis.

#### 2.9.3. MS analysis

Samples were resuspended in 0.1%TFA and then analyzed using an Ultimate 3000 nano-RSLC (Thermo Scientific, San Jose California) coupled in line with a quadrupole - orbitrap Exploris 480 via a nano-electrospray ionization source (Thermo Scientific, San Jose California) and the FAIMS pro interface. Less than 200 ng tryptic peptides were loaded on a C18 Acclaim PepMap100 pre-column (300 µm ID x 5 mm, 5 µm, 100Å, Thermo Fisher Scientific) for 1 minutes at 15 µL/min with 2% ACN, 0.1% FA in H2O and then separated on a C18 PepMap nano-column (75 µm ID x 15 cm, 3 µm, 150Å, Thermo Fisher Scientific) with a 40 minutes linear gradient from 5% to 35% buffer B (A: 0.1% FA in H2O; B: 0.1% FA in 80% ACN) followed by a regeneration step at 90% B and a equilibration at 5% B. The total chromatography was 60 minutes at 450 nl/min and the oven set to 50°C. The mass spectrometer was operated in positive ionization mode in Data-Dependent Acquisition (DDA) with one FAIMS compensation voltage (CV = -45V). The DDA cycle consisted of one survey scan or MS1 (330-1400 m/z, 90,000 FWHM) followed by MS2 spectra (HCD; 30% normalized energy; 8 m/z window; Scan Range 140-1400 m/z; 22,500 FWMH). The cycle time was limited at 1.5 sec. The Automatic Gain Control (AGC target) were set to 3E6 and 1E5 respectively for MS1 and MS2 scans, and the maximum injection time (IT) was set to 50 ms for both scan modes. Unassigned and single charged states were rejected. The exclusion duration was set for 30s with mass width was ± 10 ppm.

#### 2.9.4. MS data processing

Proteins were identified with Proteome Discoverer 2.5 software (Thermo Fisher Scientific) and Homo Sapiens proteome database (Swissprot, reviewed, release 2025_04_18 with 20644 entries). Precursor and fragment mass tolerances were set at 10 ppm and 0.02 Da respectively, and up to 2 missed cleavages were allowed. Phosphorylation (S, T, Y), Oxidation (M) and Acetyl (Protein N-term) were set as variable modification, and Carbamidomethylation (C) as fixed modification. Peptides and proteins were filtered with a false discovery rate (FDR) at 1%, rank 1. Proteins were quantified with a minimum of 1 unique peptide based on the XIC (sum of the Extracted Ion Chromatogram). The quantification values were exported in Perseus for statistical analysis involving a log[2] transform, imputation, normalization.

#### 2.9.5. Bioinformatic treatment of proteomic data

Quantification values were exported to the R environment (version 4.5.2) for statistical analysis using the DEP2 package ^6^, which includes data filtering, variance normalization and imputation of missing values prior to differential analysis. A linear model was applied to identify differentially enriched / expressed proteins. Proteins with an excessive number of missing values were excluded. Only proteins detected in at least 80% of samples were then background-corrected and normalized by variance-stabilizing transformation. Remaining missing values were imputed using the MinProb method. Principal component analysis was performed using FactoMineR and factoextra packages ^7^. For differential expression analysis, protein-wise linear models combined with empirical Bayes statistics were employed. Comparisons between panitumumab treated versus non-treated conditions were conducted for each U87 cell lines (expressing EGFR wild-type or vIII mutant) and for EVs-derived from these cells. Proteins were considered differentially detected if they met the following criteria: absolute log2 fold change ≥ 1 and adjusted p-value ≤ 0.5 (corrected by Benjamini-Hochberg multiple testing method). The results of the differential analysis were visualized using volcano plots (EnhancedVolcano package) ^8^ and hierarchical clustering analysis (ComplexHeatmap package) ^9^. To further explore proteins functions related to levels changes according panitumumab treatment, we conducted Over-Representation Analysis (ORA) from lists of differentially expressed proteins (DEPs) using functional terms associated to Gene Ontology, KEGG and Reactome using DAVID online tool ^10,11^ and only retained entries with p-adjusted inferior to 0.05 using the Benjamini-Hochberg correction. In addition, Gene Set Enrichment Analysis (GSEA) ^12^ was conducted using the ranked lists of quantified proteins based on fold change, to identify pathways exhibiting coordinated changes in protein abundance, with fgsea R package ^13^. Protein sets were obtained from the Molecular Signatures Database (MSigDB) ^14^, including the Hallmark (H) ^15^, KEGG and Reactome (C2) collections.

### 2.10. Statistical analysis

Statistical analyses were performed using GraphPad Prism 9.0. For comparisons between two normally distributed groups, Student’s t-test was used, whereas one-way or two-way analysis of variance (ANOVA) was performed for multiple-group comparisons. Unless otherwise indicated, all experiments were performed using at least three independent biological replicates. Where applicable, data are presented as mean ± standard deviation (SD). Differences were considered statistically significant at p < 0.05 (* p ≤ 0.05; ** p ≤ 0.01; *** p ≤ 0.001; **** p ≤ 0.0001).

## 3. Results

### 3.1. Validation of the U87-EGFRvIII cellular model and characterization of extracellular vesicles

Before studying the effects of panitumumab treatment, the cell lines and extracellular vesicles (EVs) were first characterized and validated. The two glioblastoma cell lines, U87 and U87-EGFRvIII, were evaluated to confirm the expected cellular phenotype and, in particular, the presence of EGFRvIII. At the same time, EVs isolated from these two cell lines were characterized to validate the isolation method and determine whether EGFRvIII was present in the EV population.

Nanoparticle tracking analysis (NTA) was first performed to characterize the size distribution and concentration of EVs isolated from U87 and U87-EGFRvIII cell cultures. The mean EV sizes in both cell lines were very closed, one was 121.1 nm for U87-derived EVs and 135.2 nm for U87-EGFRvIII-derived EVs (**Figure 1A**). The mean EV concentrations were also very similar with 5.48 × 10¹⁰ particles/mL for U87-derived EVs and 2.99 × 10¹⁰ particles/mL for U87-EGFRvIII-derived EVs. The values obtained correspond to the size range generally reported for extracellular vesicles. The expression of EV-associated proteins was assessed by Western blot in both cellular lysates and EV samples (**Figure 1B**). The tetraspanins CD81 and CD9, as well as TSG101, a protein associated with the endosomal pathway and EV biogenesis, were detected in EV samples from both cell lines, whereas calnexin, used as a negative control, was detected only in cellular lysates. No significant differences in the levels of CD81, CD9, or TSG101 were observed between EVs derived from U87 and U87-EGFRvIII cells. To further confirm the presence of EV-associated markers at the vesicle surface, CD63 expression was investigated by flow cytometry (**Figure 1C**). EVs were labeled with PKH67 and captured using magnetic beads functionalized with an anti-CD63 antibody, allowing the detection of CD63-positive EV-associated signal. More than 95% of EVs from both U87 and U87-EGFRvIII cell cultures were positive for CD63. No significant difference was observed between the two cell lines. These results further support the successful isolation of EVs and the presence of CD63 at their surface. To validate the U87-EGFRvIII cellular model, which is expected to overexpress EGFRvIII, EGFRvIII protein levels were assessed by Western blot in both cellular lysates and EV samples (**Figure 1D**). EGFRvIII was detected in U87-EGFRvIII cells as well as in the EVs derived from these cells, whereas no detectable EGFRvIII signal was observed in the control U87 cell line. The differences between U87 and U87-EGFRvIII cells were statistically significant (p < 0.05, *), while differences between EVs derived from the two cell lines reached a higher level of statistical significance (p < 0.01, **). To further investigate the presence of EGFRvIII at the surface of EVs, flow cytometry analysis was performed using PKH67-labeled EVs captured with magnetic beads (**Figure 1E**). EGFRvIII was detected on more than 90% of EVs derived from the U87-EGFRvIII cell line. This proportion was significantly higher than that observed for EVs derived from the control U87 cell line, for which no detectable EGFRvIII-associated signal was observed. These results confirm the presence of EGFRvIII at the surface of EVs released by U87-EGFRvIII cells. Overall, these results validated the U87-EGFRvIII cellular model and confirmed the successful isolation and characterization of EV derived from both cell lines. It is important to note that EGFRvIII was detected both in U87-EGFRvIII cells and in the EVs derived from them, including on their surface, which confirms the relevance of this model for the study of EGFRvIII-associated EV.

**Figure 1:**
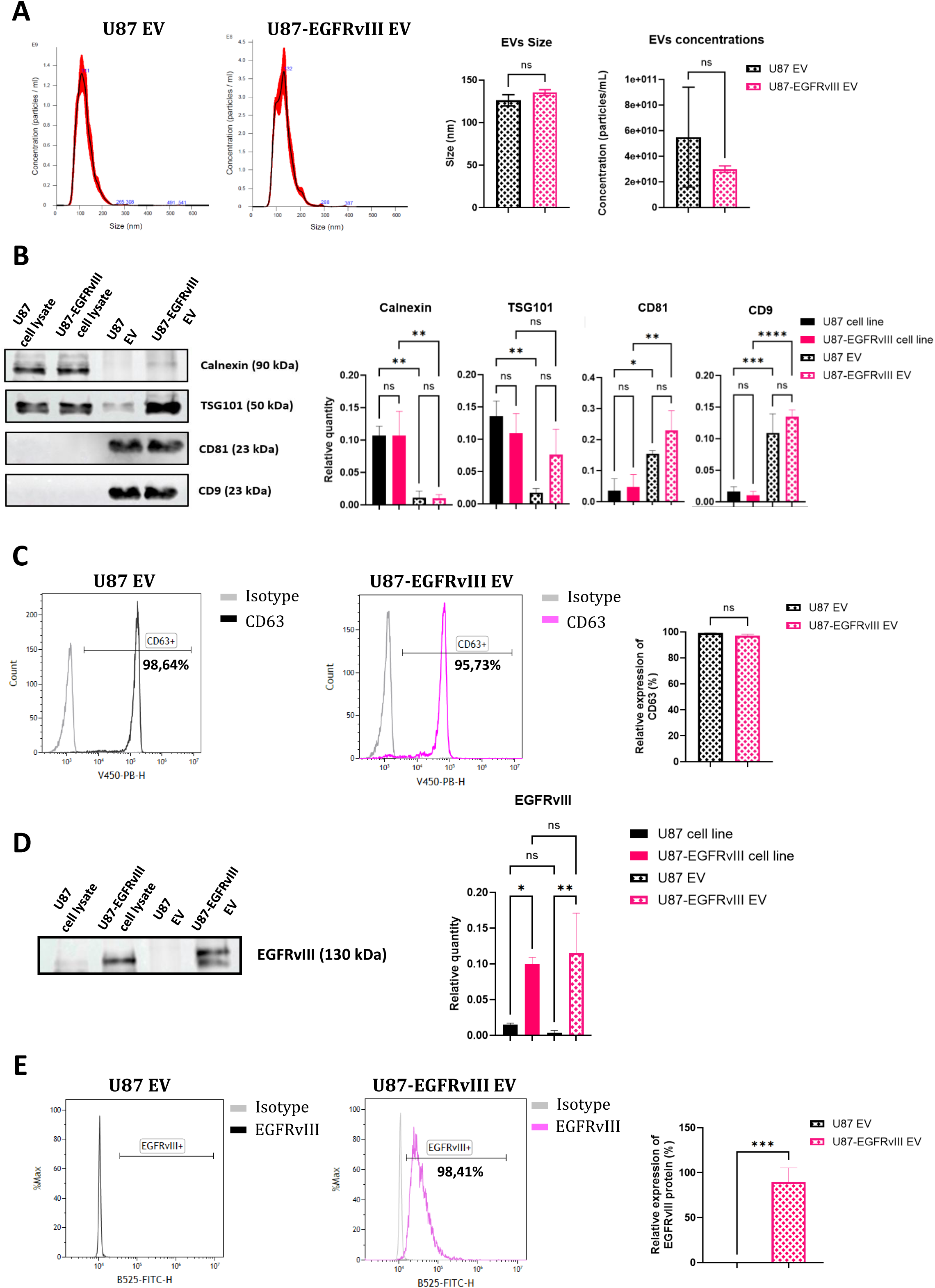
Characterization and validation of cellular models and extracellular vesicle isolation. **A**) Representative nanoparticle tracking analysis (NTA) profiles of EVs isolated from U87 and U87-EGFRvIII cell cultures (left panels). The black line represents the mean particle size distribution, and the shaded area represents the standard error of five recordings. Quantification of EV size and concentration for each cell line is shown in the right panels. The mean EV size was 121.1 nm for U87-derived EVs and 135.2 nm for U87-EGFRvIII-derived EVs, while the mean EV concentration was 5.48 × 10¹⁰ particles/mL for U87-derived EVs and 2.99 × 10¹⁰ particles/mL for U87-EGFRvIII-derived EVs. **B**) The expression of the indicated proteins was assessed by Western blot analysis in cells and extracellular vesicles from both cell lines (left panel). The corresponding protein expression quantifications are shown in the right panels. **C**) Representative flow cytometry profiles showing CD63 detection on PKH67-labeled EVs isolated from U87 and U87-EGFRvIII cell cultures (left panels). The corresponding quantification of the CD63-positive EV-associated signal is shown in the right panel. PKH67 labeling controls are not shown. **D**) EGFRvIII expression was assessed by Western blot analysis in cells and extracellular vesicles from both cell lines (left panel). The corresponding quantification of EGFRvIII expression is shown in the right panel. **E**) Representative flow cytometry profiles showing EGFRvIII detection on the surface of PKH67-labeled EVs isolated from U87 and U87-EGFRvIII cell cultures (left panels). The corresponding quantification of the EGFRvIII-positive EV-associated signal is shown in the right panel. PKH67 labeling controls are not shown. Performed in biological triplicate. * p < 0.05; ** p < 0.01; *** p < 0.001, **** p < 0.0001. ns: no significance.

### 3.2. Effect of panitumumab treatment on cellular and extracellular vesicle-associated EGFRvIII

After the validation of the cellular models and the isolation of EVs, the effect of panitumumab treatment was investigated in U87 and U87-EGFRvIII cells. Cells were cultured for 72 h in the presence of 10 µg/mL panitumumab, after which both cells and supernatants were collected. Cell lysates and EVs isolated from the corresponding conditioned culture media were then analyzed. The goal was to evaluate the effects of panitumumab on the expression of EGFR-related proteins and on the characteristics of the EVs, including their size, concentration, and the impact on EGFRvIII on their surface.

To investigate the effect of panitumumab treatment on EGFR-related proteins, Western blot analyses were performed on cellular lysates and corresponding EV samples from U87-MG and U87-EGFRvIII cells (**Figure 2A**). Calnexin was detected only in cellular lysates and the EV-associated markers CD81 and CD9 were detected in EV samples from both cell lines. The levels of CD81 and CD9 did not significantly differ between untreated and treated conditions in either cell line, indicating that panitumumab treatment did not significantly affect the levels of these EV-associated markers **(Supplementary Figure S1)**. EGFR was detected in U87-EGFRvIII cells and their corresponding EVs, with a non-significant decrease in cellular lysates following treatment, while a significant reduction was observed in EVs (** p < 0.01). Regarding EGFR-related proteins, phosphorylated EGFR (p-EGFR) was detected exclusively in U87-EGFRvIII cells and their corresponding EVs. Following panitumumab treatment, p-EGFR levels were significantly reduced in EVs (* p < 0.05), whereas the decrease observed in cellular lysates did not reach statistical significance. In the same way, Finally, EGFRvIII was detected exclusively in U87-EGFRvIII cells and their derived EVs. A highly significant decrease in EGFRvIII levels was observed in EVs following treatment (**** p < 0.0001), whereas no statistically significant difference was observed in cellular lysates. Overall, these results showed that panitumumab treatment appeared to have more pronounced effect on EGFR-related proteins in EVs than in the corresponding cellular lysates, particularly for EGFRvIII. To determine whether panitumumab treatment affected EV characteristics, the mean EV size and particle concentration were assessed by nanoparticle tracking analysis (NTA) (**Figure 2B–C**). No significant differences were observed in either the mean EV size or particle concentration following panitumumab treatment in both cell lines. Finally, EGFRvIII at the EV surface was assessed by flow cytometry after treatment with panitumumab. (**Figure 2D**). In untreated conditions (NT), EGFRvIII was detected at the surface of approximately 80% of EVs derived from U87-EGFRvIII cells. Following panitumumab treatment (T), no detectable EGFRvIII-associated signal was observed at the EV surface, corresponding to a significant decrease compared with untreated conditions (** p < 0.01). As expected, no EGFRvIII-associated signal was detected at the surface of EVs derived from U87 cells, either before or after treatment.

**Figure 2:**
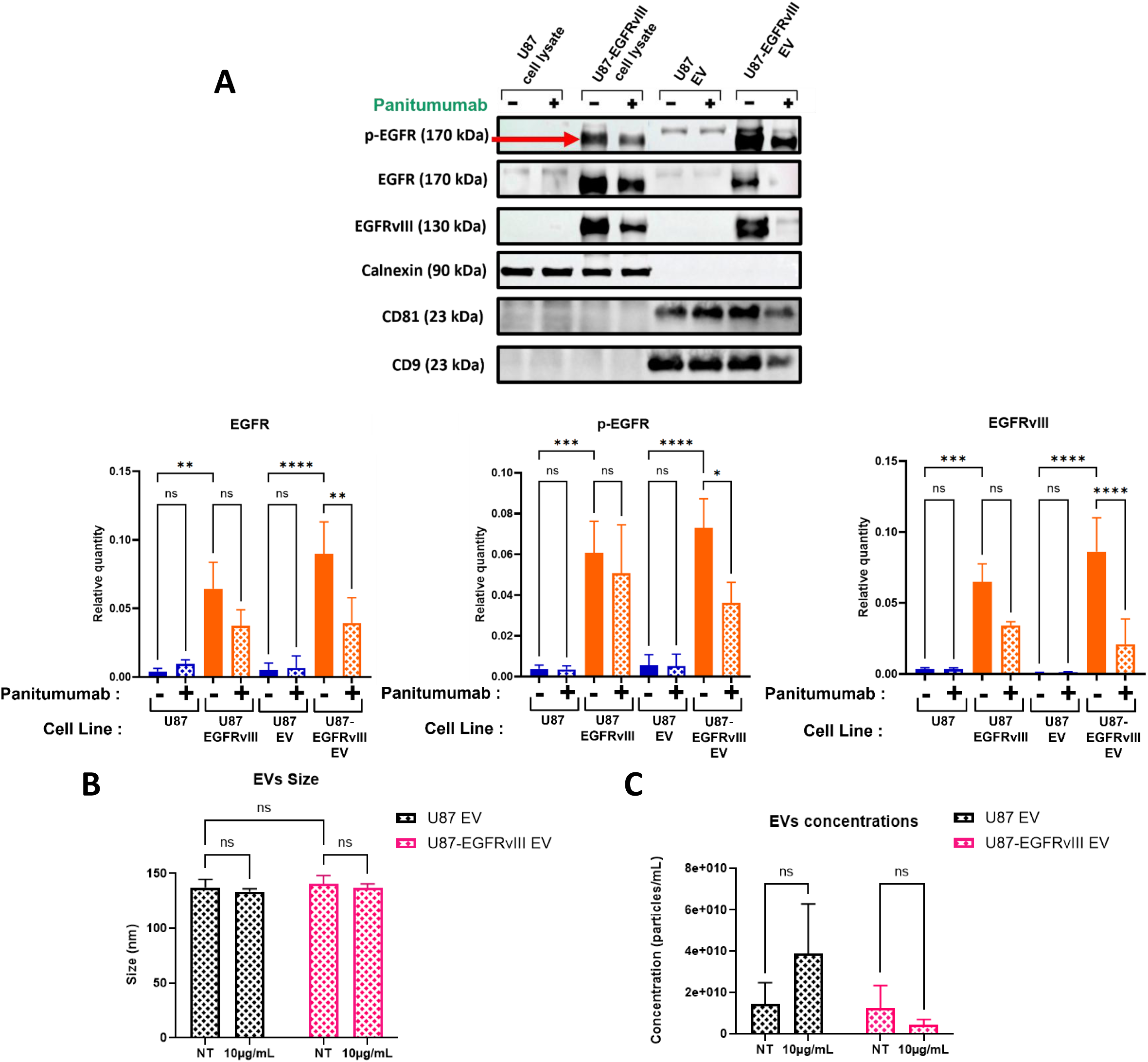
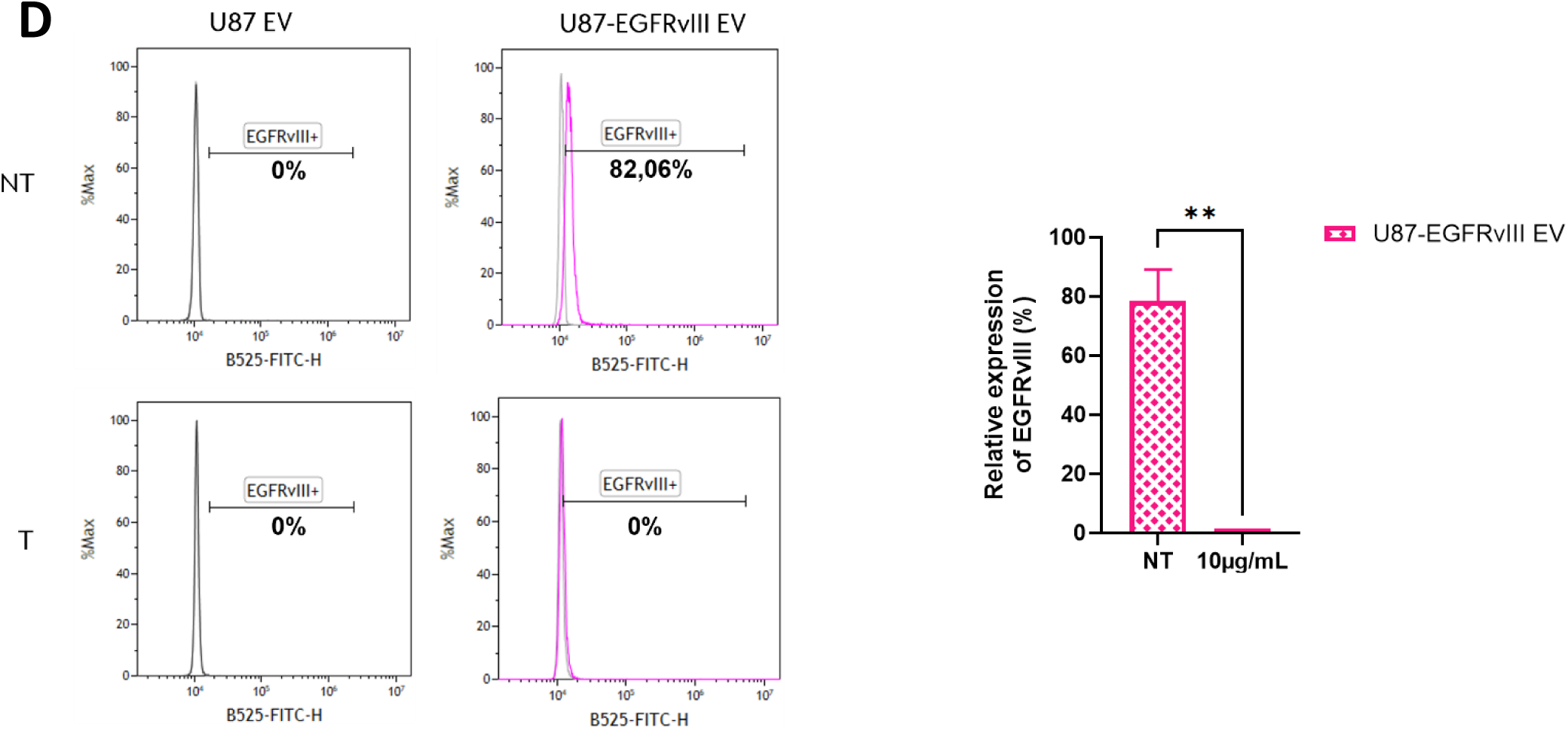
Effect of panitumumab treatment on cellular and extracellular vesicle-associated protein expression and EV characteristics. **A**) Western blot analysis of EGFR, phosphorylated EGFR (p-EGFR), EGFRvIII, and other proteins in U87 and U87-EGFRvIII cells and their corresponding EVs following panitumumab treatment (upper panel). The corresponding quantification of EGFR, p-EGFR, and EGFRvIII protein levels is shown in the lower panels for cells and EVs. **B**) Quantification of the mean EV size determined by nanoparticle tracking analysis (NTA) in U87 and U87-EGFRvIII cell cultures following panitumumab treatment. The mean EV size was 137.13 nm for non-treated (NT) U87-derived EVs, 133.3 nm for treated (T) U87-derived EVs, 140.5 nm for non-treated U87-EGFRvIII-derived EVs, and 136.5 nm for treated U87-EGFRvIII-derived EVs. **C**) Quantification of EV particle concentration determined by nanoparticle tracking analysis (NTA) in U87 and U87-EGFRvIII cell cultures following panitumumab treatment. The mean EV concentration was 1.43 × 10¹⁰ particles/mL for non-treated (NT) U87-derived EVs, 3.8 × 10¹⁰ particles/mL for treated (T) U87-derived EVs, 1.2 × 10¹⁰ particles/mL for non-treated U87-EGFRvIII-derived EVs, and 4.38 × 10¹⁰ particles/mL for treated U87-EGFRvIII-derived EVs. **D**) Representative flow cytometry profiles showing the detection of EGFRvIII on the surface of PKH67-labeled EVs isolated from U87 and U87-EGFRvIII cell cultures, with or without panitumumab treatment (left panels). The corresponding quantification of the EGFRvIII-positive EV-associated signal is shown in the right panel. PKH67 labeling controls are not shown. Performed in biological triplicate. * p < 0.05; ** p < 0.01; *** p < 0.001, **** p < 0.0001. ns: no significance.

### 3.3. Effect of panitumumab treatment on the proteomic profiles of cells and extracellular vesicles

To further investigates the impact of panitumumab, principal component analysis (PCA) was performed on the proteomic profiles of cellular and extracellular vesicles lysates derived from U87 and U87-EGFRvIII cells untreated (NT) or treated (T) with panitumumab (**Figure 3**). Each PCA was conducted using normalized protein abundance levels for proteins detected in at least 80% of the samples. This screening process resulted in 2 877 proteins for U87 cells, 2 798 for U87-EGFRvIII cells, 733 for U87-derived EVs, and 862 for U87-EGFRvIII-derived EVs.

**Figure 3:**
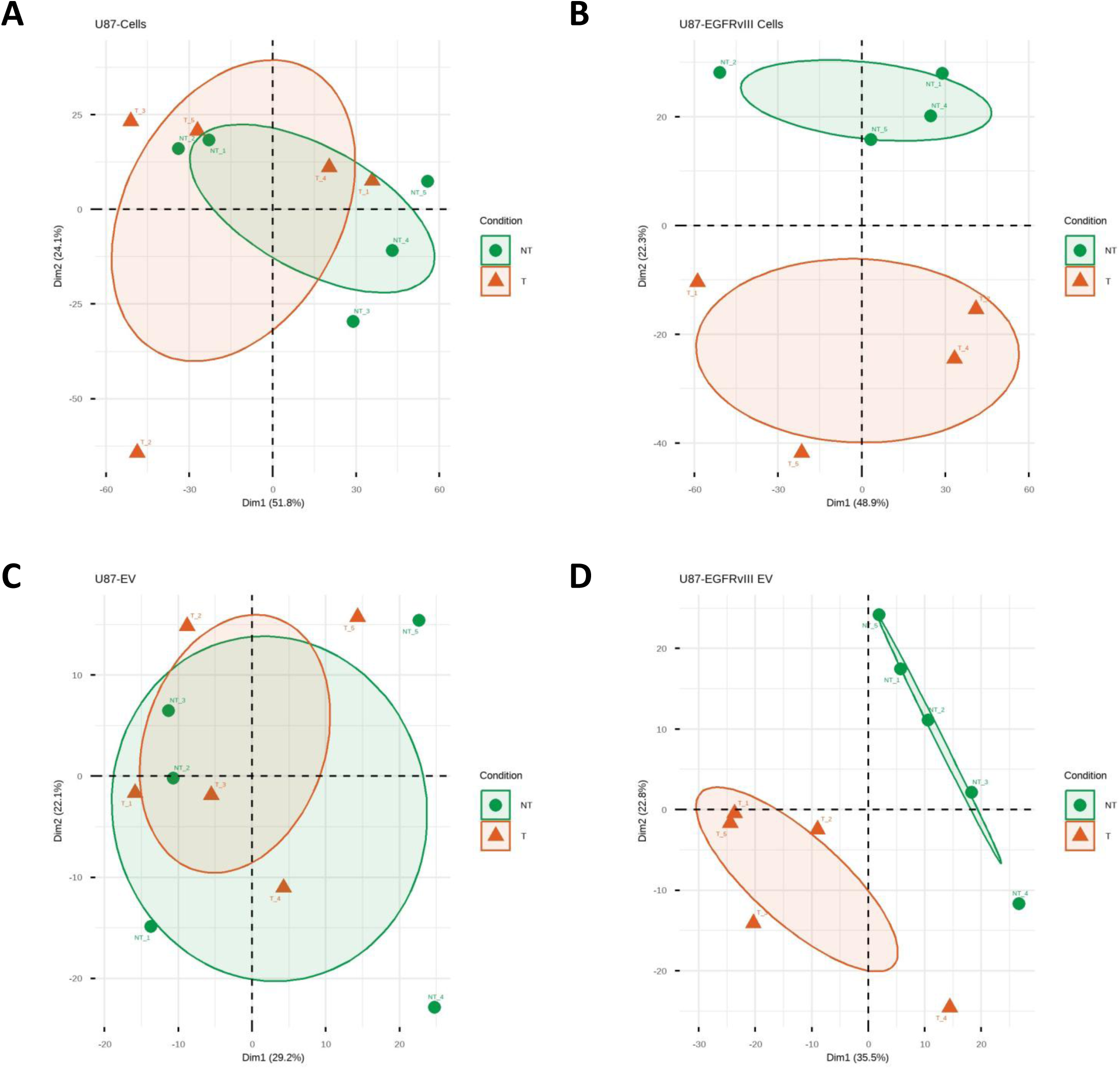
Proteomic profiling of U87 cells or EV-derived from U87 cells harboring EGFR wild type or vIII mutant, treated or not with panitumumab. Principal component analysis of proteomic data in U87 cells (**A**), U87-EGFRvIII cells (**B**), EVs derived from U87 cells (**C**) and EVs derived from U87-EGFRvIII cells (**D**) treated or not by panitumumab reveals a clear distinction between no-treated and treated samples, only for EGFRvIII-expressed cells or EVs. Samples were obtained from four (U87-EGFRvIII cells) or five (U87 cells, U87 EVs and U87-EGFRvIII EVs) independent experiments. The absence of overlap between the confidence ellipses of the barycenters of each group indicates a distinct proteomic profile. Each point represents one sample, colored according to culture condition and treatment (No treated, NT: green, Treated, T: orange). Each PCA was conducted from normalized levels of proteins detected in 80% of samples (U87 cells: 2877; U87-EGFRvIII cells: 2798, U87 EVs: 733 and U87-EGFRvIII EVs: 862 proteins).

Our results demonstrate that the proteomic profile of U87-treated cells is closely similar to parental U87 cells, whereas U87-EGFRvIII-treated cells exhibit a distinct profile compared to their untreated counterparts (**Figure 3A and 3B**). These results suggest that panitumumab has no real effect on U87 cells in which EGFR expression is limited and EGFRvIII is absent compared to U87-EGFRvIII cells. In the latter, EGFRvIII and EGFR expression are significantly higher, explaining their more pronounced response to treatment compared to the parental cells. Similar results were observed with proteomic profiles derived from EVs. As observed previously, U87-derived EVs shows an overlap between the convex hull of the untreated and treated samples (**Figure 3C**). On the contrary, the proteomic profile of EVs derived from treated EGFRvIII-cells appeared distinct from that of EVs from non-treated cells, as showed by principal component analysis (PCA), in which two groups are clearly distinguished (**Figure 3D**). This marked separation in the convex hulls on the PCA between EVs from treated and non-treated cells suggests that the treatment has an indirect influence on the extracellular vesicle cargo and EV biogenesis pathway.

Overall, the PCA analysis revealed different patterns of proteomic variation following panitumumab treatment in the two cellular models. While the profiles of U87 cells and their derived EVs show no clear differences between untreated and treated conditions, distinct profiles are observed between untreated and treated U87-EGFRvIII cells and their respective EVs compared to their native cells and corresponding EVs. Those results suggest that the proteomic response to panitumumab might be different according to the EGFRvIII status of the cellular model and highlight the role of EV proteomic profiles to reflect changes related to treatment.

### 3.4. Identification of panitumumab-associated proteomic changes in U87-EGFRvIII cells and their extracellular vesicles

Based on the PCA, which revealed a clearer separation between proteomic profils of untreated and treated conditions in U87-EGFRvIII cells and their EVs, we chose to focus on these two types of samples to further investigate the proteomic changes induced by the panitumumab treatment. Thus, a differential proteomic analysis was performed to compare “treated” and “untreated” conditions using U87 cells expressing wild-type or mutant (vIII) EGFR, along with the EVs isolated from both cell types before and after panitumumab treatment. The differential expression analysis performed on these samples and the results were represented using volcano plots and heatmaps in order to identify Differentially Expressed Proteins (DEPs) and to compare their abundance patterns between untreated (NT) and treated (T) conditions (**Figure 4**).

**Figure 4:**
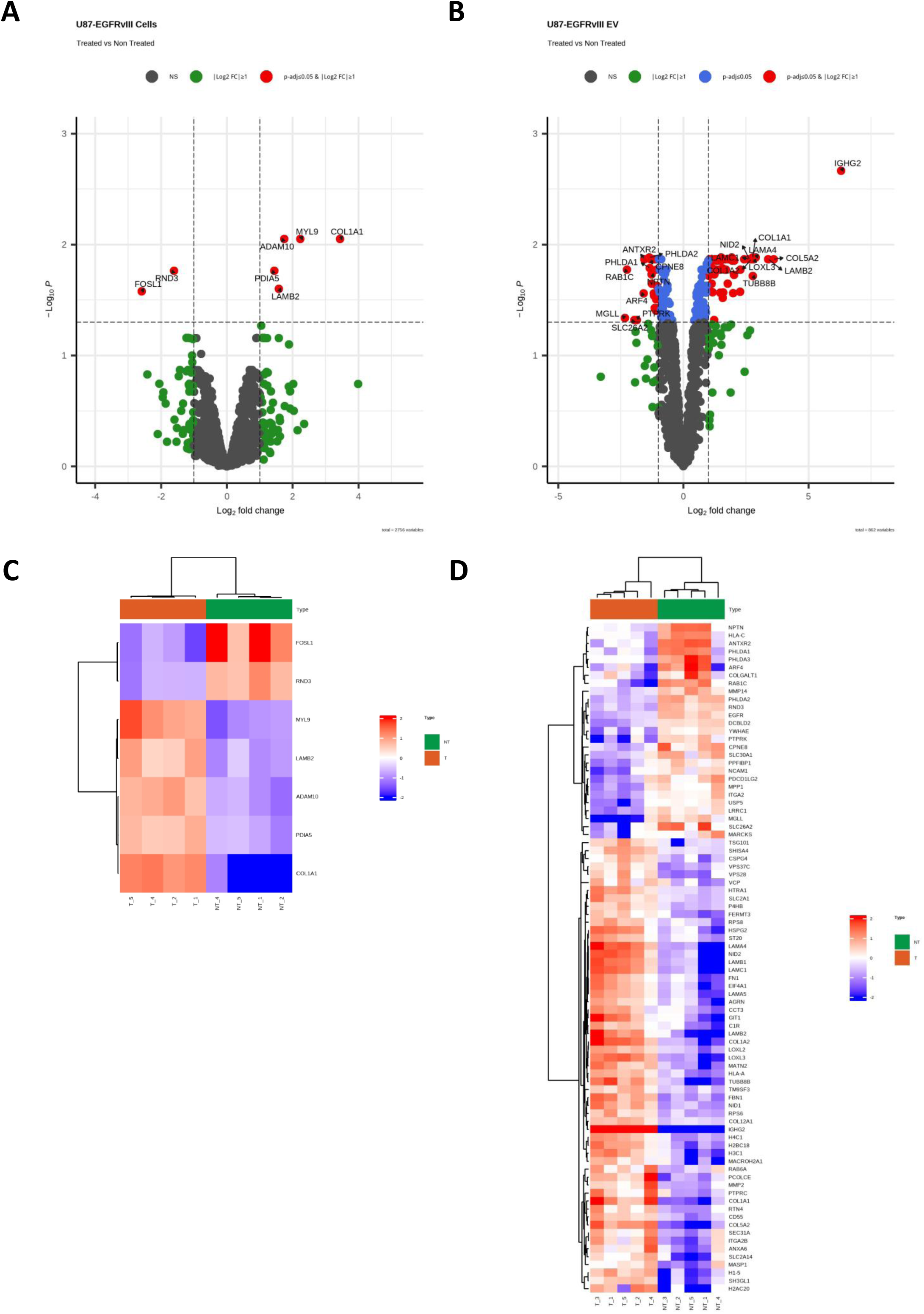
Proteomic signature associated to panitumumab treatment on U87-EGFRvIII cells and their extracellular vesicles (EVs). Differential proteome analysis of panitumumab treated as compared to no treated conditions were conducted from U87 cells expressing EGFR wt or mutant (vIII) and from EVs produced by these two types of cells. **(A and B)** Results obtained from U87-EGFRvIII cells (**A**) and from U87-EGFRvIII EVs (**B**) were displayed by volcano plot. Each dot represents one protein and selected differentially expressed proteins (DEPs) according to threshold criteria (dashed lines; adjusted P value ≤ 0.05 and absolute log2 fold change ≥ 1) appeared in red. Among 7 DEPs in U87-EGFRvIII cells, 5 were up-regulated and 2 down-regulated in treated as compared to no treated conditions. Among 84 DEPs in U87-EGFRvIII EVs, 57 were up-regulated and 27 down-regulated in treated as compared to no treated conditions. Labels corresponds to most up- and down-deregulated proteins. **(C and D)** Heatmap displaying the level of deregulation of DEP between treated as compared to non-treated U87-EGFRvIII cells **(C)** and between their respective EVs **(D)**. Protein levels were log transformed and median-centered and are displayed on heatmap with color gradient from blue to red, reflecting low to high expression level, respectively. Samples (columns) are hierarchically clustered using Pearson correlation distance and average linkage method.

In U87-EGFRvIII cells, only seven differentially expressed proteins were identified with an adjusted p-value ≤ 0.05 and an absolute log₂ fold change ≥ 1 (**Figure 4A**). Among those seven DEPs, five were up-regulated MYL9, COL1A1, ADAM10, LAMB2 and PDIA5 and two were down-regulated FOSL1 and RND3, following panitumumab treatment. The corresponding heatmap shows the relative abundance of these seven DEPs in untreated and panitumumab-treated U87-EGFRvIII cells, illustrating the differences in protein expression identified by the differential expression analysis (**Figure 4C**).

Regarding the results obtained with U87-EGFRvIII-derived EVs, we could see a more extensive set of changes (**Figure 4B**). Indeed, a total of 84 proteins were differentially expressed, including 57 upregulated and 27 downregulated proteins following panitumumab treatment. Among the proteins showing an increase abundance, we can find COL1A1, COL5A2 and LAMB2. Notably, COL1A1 and COL5A2 have previously been detected in both cancer-associated fibroblasts and their derived exosomes, supporting the association of extracellular matrix proteins with EVs in cancer models ^16^. On the other hand, downregulated proteins include ANTXR2, PHLDA1, PHLDA2, RAB1C and MGLL. The upregulation of COL1A1, COL5A2 and LAMB2 highlights changes in EV-associated proteins involved in extracellular matrix organization and cell–matrix interactions ^16,17^. The heatmap also demonstrates differences in protein abundance with distinct clustering patterns between untreated and treated EV samples (**Figure 4D**). In order to further investigate the relationship between the 84 DEPs proteins identified, a protein-protein interaction network was generated using STRING database (**Supplementary Figure S2**). It appears that of the 84 proteins, 73 were connected to at least one other protein, resulting a formation of 307 edges. This analysis identified the known and predicted interactions between the different proteins and completed the differential proteomic analysis.

Overall, differential proteomic analysis identified a higher number of differentially expressed proteins in U87-EGFRvIII-derived EVs than in their corresponding cellular lysates following panitumumab treatment. Altogether these results suggest that the effect of panitumumab on U87-EGFRvIII cells was closely linked to changes in the protein composition of EVs released by U87-EGFRvIII cells. Further investigation of these proteins could help identify biological pathways involved in these changes and provide insights into the mechanisms underlying EGFRvIII regulation, trafficking, and fate following panitumumab treatment.

### 3.5. Functional enrichment analysis of differentially detected proteins in U87-EGFRvIII-derived EVs following panitumumab treatment

To further investigate the biological process associated with the proteomic changes observed in U87-EGFRvIII-derived EVs, a functional enrichment analysis was performed using the DAVID online tool on the 84 differentially expressed proteins (DEPs) detected in U87-EGFRvIII EVs from panitumumab-treated and untreated conditions (**Figure 5A**). Gene ontology (GO) analysis highlighted several terms associated with the extracellular matrix (ECM) organization, notably ECM structural constituents, basement membrane, extracellular matrix, and the solid phase of the interstitial matrix. The Reactome pathway analysis also identified several enriched pathways as significant such as ECM organization, ECM proteoglycans, and collagen formation. Thus, we can find in these categories different proteins involved in ECM structure and organization, such as collagens, laminins, fibronectin, and other matrix-associated proteins. Furthermore, an additional enriched pathways related to the epigenetic regulation of gene expression by MLL3/MLL4 complexes has been highlighted. The GO terms and Reactome pathways identified by the enrichment analysis, together with their associated proteins, are detailed in **Supplementary Table S1**.

**Figure 5:**
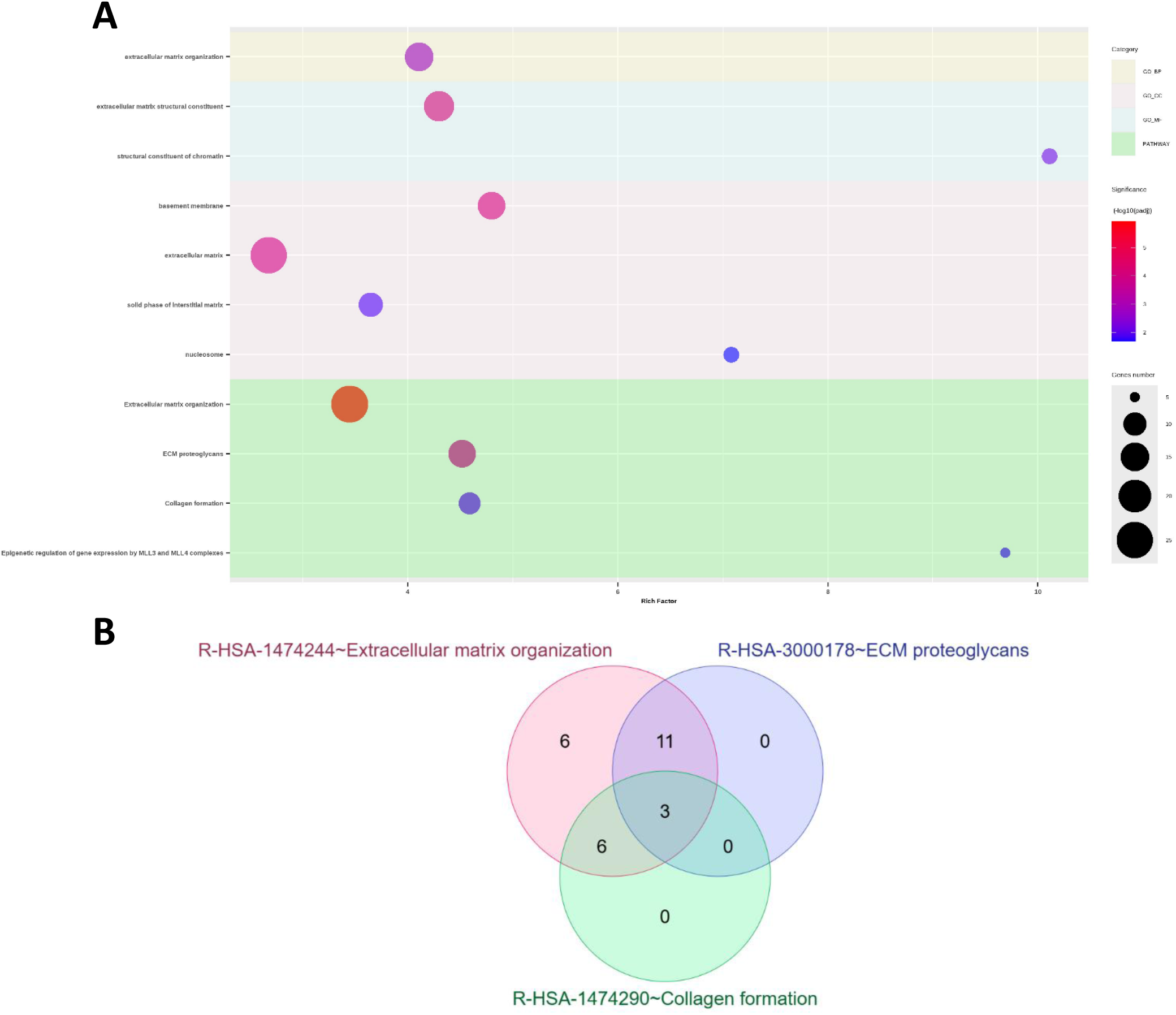
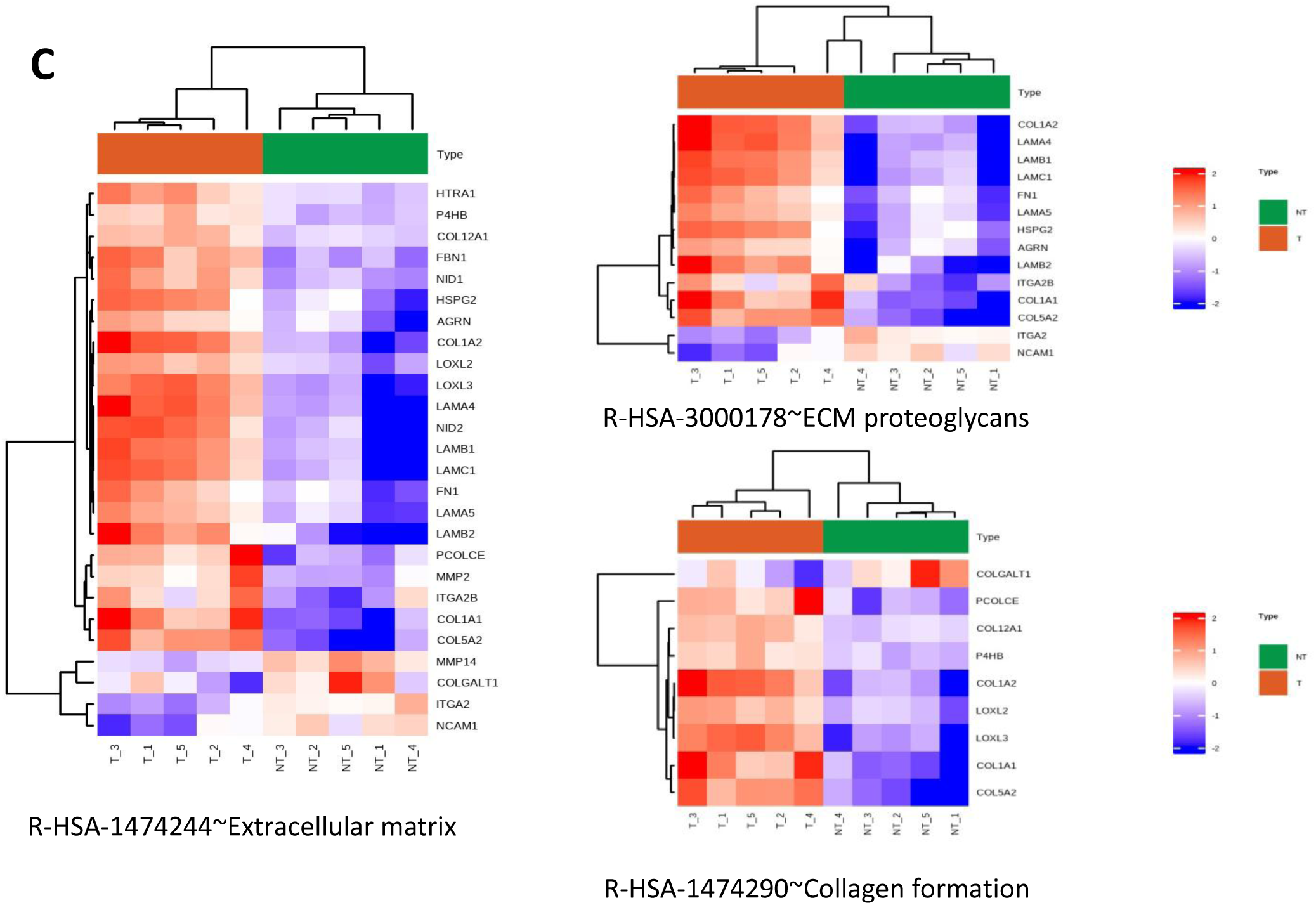
Functional enrichment analysis of differentially detected proteins in U87-EGFRvIII-derived EVs following panitumumab treatment. **A)** Gene Ontology (GO) terms and Reactome pathways enriched among the 84 differentially detected proteins identified in EVs from untreated and panitumumab-treated U87-EGFRvIII cells, using the DAVID online tool. **B)** Venn diagram showing the overlap between proteins associated with the three enriched Reactome pathways: ECM organization, ECM proteoglycans, and collagen formation. **C)** Hierarchical clustering heatmaps showing the relative abundance patterns of proteins associated with each of these pathways in untreated and panitumumab-treated EV samples.

In order to investigate the relationships between the differentially expressed proteins, a protein–protein interaction network was generated using the STRING database (**Supplementary Figure S2**). Of the 84 DEPs, 73 were connected to at least one other protein, forming a network comprising 307 edges. This network illustrates the connectivity between proteins affected by panitumumab treatment, although the predicted interactions do not necessarily indicate direct physical interactions or establish functional relationships in the experimental model.

A Venn diagram was then used to observe the overlap between the three main Reactome pathways (**Figure 5B**). This analysis showed eleven proteins shared between the ECM organization and ECM proteoglycans pathways, while six proteins were found in common between ECM organization and collagen formation. In total, three proteins were common to all three pathways, demonstrating an overlap between these related biological processes.

After that, hierarchical clustering heatmaps were generated to further investigate the abundance patterns of proteins associated with ECM organization, ECM proteoglycans, and collagen formation pathways (**Figure 5C**). The heatmaps showed different proteins abundance patterns between untreated and treated EV samples. Several proteins including collagens, laminins, and other ECM-associated components were found in all three pathways. It was also possible to identify COL1A2, COL1A1 and COL5A2 as the proteins commonly found in all three pathways. Although some differences were found across individual samples, the overall abundance patterns indicated changes in the EV proteome due to panitumumab treatment.

Overall, functional enrichment and hierarchical clustering heatmaps analysis identified ECM organization, ECM proteoglycans, and collagen formation as the main pathways associated with proteomic changes in U87-EGFRvIII EVs treated with panitumumab. These results suggest that panitumumab induces changes in EVs proteins that can be involved in microenvironment remodeling.

## 4. Discussion

Glioblastoma (GB) is one of the most common and aggressive brain tumors, also characterized by frequent relapse after treatment. EGFRvIII, a truncated variant of EGFR, is found in approximately 40% of GBs and has been detected in extracellular vesicles (EVs) derived from glioma cells ^3,4^. In this context, EVs emerge as potential sources of biomarkers, as they can be found in the bloodstream and reflect potential changes in their cells of origin. Nevertheless, the effects of anti-EGFR targeted therapies on the protein composition of glioblastoma-derived EVs, particularly those expressing EGFRvIII, remain incompletely characterized. In this study, we investigated the effect of a targeted therapy directed against EGFR and EGFRvIII, panitumumab, on EGFR/EGFRvIII and compared its impact on the changes of proteomic profiles of U87 and U87-EGFRvIII cells and their respective derived EVs. Our results showed that the treatment led to a decrease in EGFR, its active form p-EGFR, and, most notably, EGFRvIII accompanied with EV proteome changes and differential protein abundance between both EV populations. The main enriched pathways were identified through a functional enrichment analysis performed on the EV proteome. Functional enrichment analysis revealed that main enriched pathways following treatment were extracellular matrix (ECM) organization, ECM proteoglycans, and collagen formation were the main enriched pathways. These results suggest that EV-derived protein profiles might provide valuable circulating biomarkers that reflect the adaptation mechanisms developed by glioblastoma cells to resist and escape treatment.

One of the key findings obtained was a significant decrease in EGFR-associated proteins, particularly EGFRvIII, in EVs derived from U87-EGFRvIII following treatment with panitumumab. In contrast, although a similar tendency was observed in U87-EGFRvIII cell lysates, the results were not statistically significant. Flow cytometry results also revealed that EGFRvIII could no longer be detected on the surface of these vesicles after treatment compared to untreated conditions. Together, these findings suggest that the effects of panitumumab on EGFRvIII may differ between cells and their derived EVs. Panitumumab binding could thus affect receptor trafficking, its degradation, or its incorporation into vesicles. Indeed, studies have shown that panitumumab induces the internalization of EGFRvIII followed by its recycling to the plasma membrane, rather than degradation of the receptor ^18^. This observation is convenient with our results. Indeed, the significant decrease in EGFRvIII observed in EVs after treatment could be due, at least in part, to a change in the receptor’s intracellular trafficking and, consequently, to the biogenesis of EVs. This would imply that the receptor avoids intracellular degradation and is instead redistributed, altering its incorporation into EVs. Further experiments using complementary approaches, including confocal microscopy to assess EGFRvIII colocalization with endosomal and lysosomal markers (Rab5, Rab11 and LAMP1), could help investigate whether changes in receptor trafficking contribute to its reduced detection in EVs. To better understand which proteins were activated by this mechanism and to characterize these proteins, we decided to study changes in the proteomic profile before and after treatment, both in the two distinct cell lines and in their respective EVs.

The changes of EGFRvIII expression in EVs are associated with significant difference in the number of differentially expressed proteins (DEPs) in cell and EVs following treatment with panitumumab. Thus, only seven DEPs were identified in U87-EGFRvIII cells, whereas 84 proteins were identified in the EVs, of which 57 were upregulated and 27 were downregulated. These results demonstrate that panitumumab treatment primarily targets EGFRvIII-expressing cells, inducing alterations in their protein expression profiles. The impact of this phenotypic shift is not restricted to the intracellular environment but extends to the secreted extracellular vesicles. Consequently, variations in the exosomal protein cargo are indirectly driven by the effects of panitumumab on the parental cells. Analyzing the proteomic content of these exosomes therefore enables the identification of key distal mechanisms activated by panitumumab-targeted tumor cells and offers new insights into signaling networks and resistance mechanisms activated by treated tumor cells.

The functional enrichment analyses provided further insight into these changes by identifying the extracellular matrix (ECM) organization, ECM proteoglycans, and collagen formation pathways as the main enriched pathways in the EVs derived from U87-EGFRvIII. Three proteins associated with these functional pathways were thus identified as common: COL1A1, COL1A2, and COL5A2. These results highlight the importance of collagen-associated proteins, in the proteomic profile of EV following treatment. Indeed, these observations are of prime interest since, in the context of glioblastoma, the ECM plays a significant role in the interaction between the tumor and the microenvironment, as well as in invasion and disease progression ^19^. Studies have shown that COL1A2 expression is increased in glioblastomas compared to healthy brain tissue, and that its inhibition reduces glioblastoma cell proliferation and invasion ^20,21^. COL1A1 shares a similar function with the COL1A2, and like the latter, it plays a pivotal role in driving the mesenchymal transition of gliomas and is a major driver of tumor aggressiveness exerting a strong regulatory control over their malignant properties ^22^. It has also been described as contributing to radiation resistance in hypoxic glioma cells ^23^. Finally, COL5A2, as well as COL1A1 and COL1A2, have been identified as being negatively correlated with overall survival in gliomas and as playing a role in the regulation of their immunosuppressive microenvironment ^24^. Furthermore, Collagen type I triggers mTOR activation independently of Akt, thereby driving resistance to EGFR-TKIs ^25^. Thus, the combination of mTOR inhibitor and EGFR-TKI could prove useful in overcoming the resistance caused by Collagen I. The presence of these proteins across related pathways suggests that the observed changes involve several components of the collagen-associated extracellular environment. These results suggest that panitumumab treatment induces changes in the structure of the ECM as well as in cell-matrix interactions. However, it is important to note that an increase abundance of these proteins in EVs does not necessarily imply an increase expression of these proteins in the origin cells.

Previous studies have already put on evidence that changes could be observed in the cargo of GB EVs following treatment. Indeed it has been shown that the temozolomide was likely to alter the composition of the EVs produced by glioblastoma stem-like cells, resulting in an enrichment of proteins associated with cell adhesion ^26^. Similarly, molecular changes induced by the development of resistance to temozolomide in GB stem cells, as well as evidence of this resistance, were observed in their EVs ^27^. Although no mechanisms of resistance to EGFR inhibitors (TKIs) have ever been observed in glioblastoma, such mechanisms have been identified in non-small-cell lung cancer (NSCLC), where resistance to osimertinib could be induced by the intercellular transfer of wild-type EGFR via exosomes ^28^. We hypothesized that the transfer of alpha-1 (COL1A1) and alpha-2 (COL1A2) chains of type I collagen could induce resistance to EGFR-TKI such panitumumab in neighboring GB cells as previously demonstrated in NSCLC with osimertinib ^25^. Thus, even though studies regarding glioblastoma mainly focus on temozolomide rather than EGFR-TKI or panitumumab, they highlight the possibility of treatment-related cellular changes that are subsequently accompanied by alterations in the cargo of EVs ^29,30^. Thus, the changes we observed in collagen-associated proteins may reflect modifications in the molecular status of U87-EGFRvIII cells following treatment.

Consequently, these results suggest that proteins associated with EVs might be potential biomarkers for monitoring treatment response in GB. Currently, monitoring the progression of GB relies on neuroimaging as well as highly invasive tumor biopsies, which are not always possible depending on the patient’s overall condition. In this context, EVs may provide a less invasive source of information on tumor status and treatment-related changes. Thus, it has been demonstrated that changes induced by surgical resection were correlated with modifications of EV concentration suggesting that EVs could provide insights about the tumor, treatment response, and disease monitoring ^31^. In our study, COL1A1, COL1A2, and COL5A2 might be potential candidates for further investigation because they were associated with all three enriched ECM-related pathways. Nevertheless, the abundance of these candidate biomarkers should first be validated using targeted proteomics or immunoassays, followed by validation in additional glioblastoma models and patient-derived samples.

In conclusion, our study revealed changes in EGFR-related proteins and in the proteomic profiles of EVs derived from U87-EGFRvIII cells following panitumumab treatment. Functional enrichment analysis highlighted pathways related to extracellular matrix (ECM) organization, with COL1A1, COL1A2, and COL5A2 shared across the three main enriched pathways. These findings highlight collagen-associated proteins as potential biomarker of treatment-related molecular changes in glioblastoma. Further studies are needed to determine whether these changes contribute to tumor progression or treatment resistance and to assess the potential of these proteins as biomarkers for monitoring treatment response in glioblastoma.

