## Supplementary Figure S1 for "Effects of panitumumab on the proteome of treated EGFRvIII glioblastoma cells and on their extracellular vesicles"

**Supplementary Figure S1: Western blot quantification of cellular and extracellular vesicle protein profiles in untreated and panitumumab-treated U87 and U87-EGFRvIII models.** Quantification of Calnexin, CD81, and CD9 protein levels is shown for cellular and EV samples. TotalStain Q was used to assess equal protein loading. Performed in biological triplicate. * $p < 0.05$; ** $p < 0.01$; *** $p < 0.001$, **** $p < 0.0001$. ns: no significance.

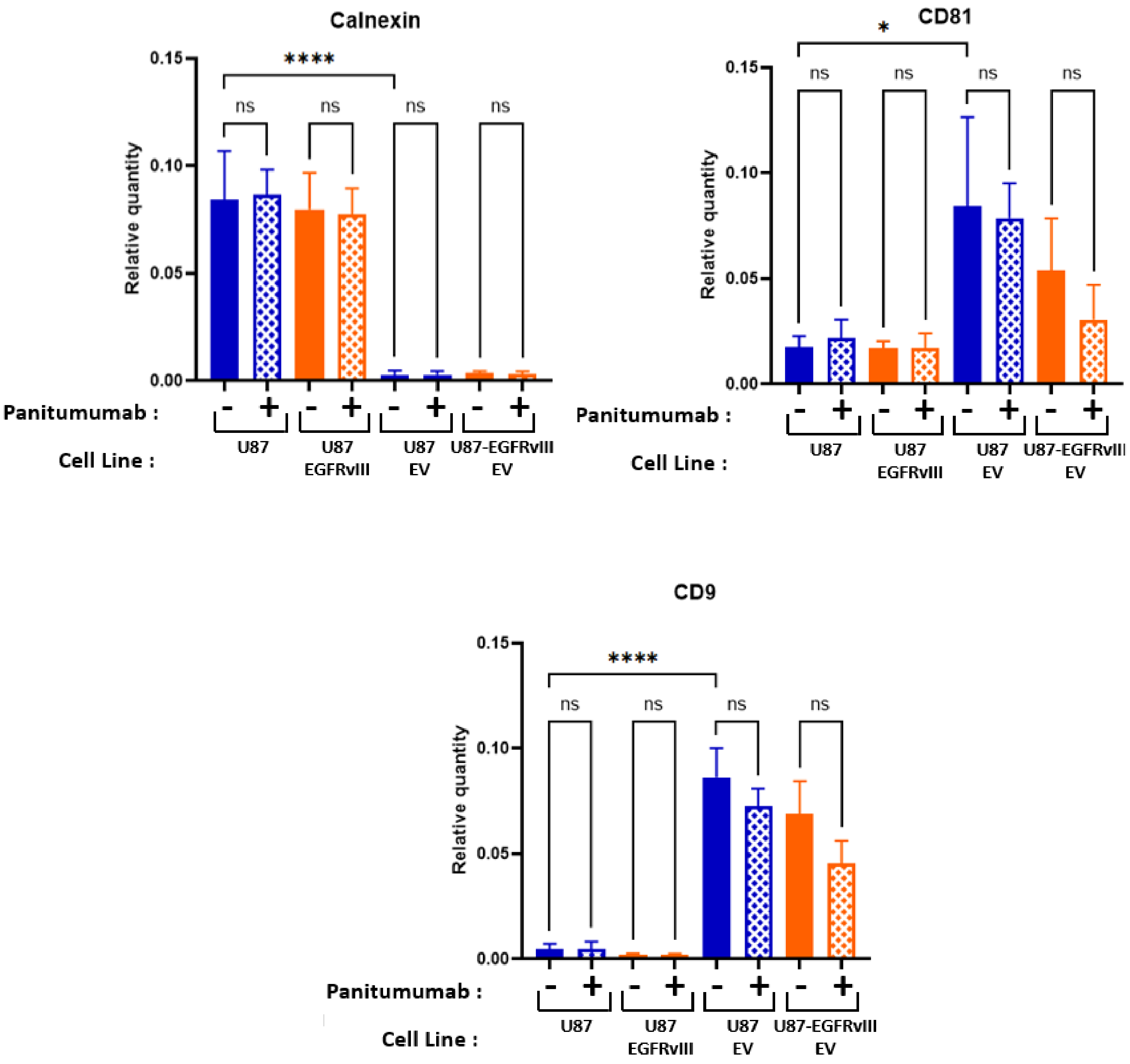
