## Supplementary Figure S2 for "Effects of panitumumab on the proteome of treated EGFRvIII glioblastoma cells and on their extracellular vesicles"

**Supplementary Figure S2: Protein–protein interaction network of differentially expressed proteins in U87-EGFRvIII-derived extracellular vesicles following panitumumab treatment.** The 84 differentially expressed proteins (DEPs) identified in U87-EGFRvIII-derived extracellular vesicles following panitumumab treatment were mapped to the STRING database to construct a protein–protein interaction network. Of these, 73 proteins were connected to at least one other protein, forming a network comprising 307 edges (interactions). Edge thickness represents the STRING confidence score (0–1), reflecting the reliability of the predicted interactions. Blue edges indicate physical interactions supported by experimental data from interactome databases. The color gradient from blue to red represents fold-change values, with blue indicating downregulated proteins and red indicating upregulated proteins following panitumumab treatment. Node size is proportional to the degree of each protein, reflecting the number of interactions within the network.

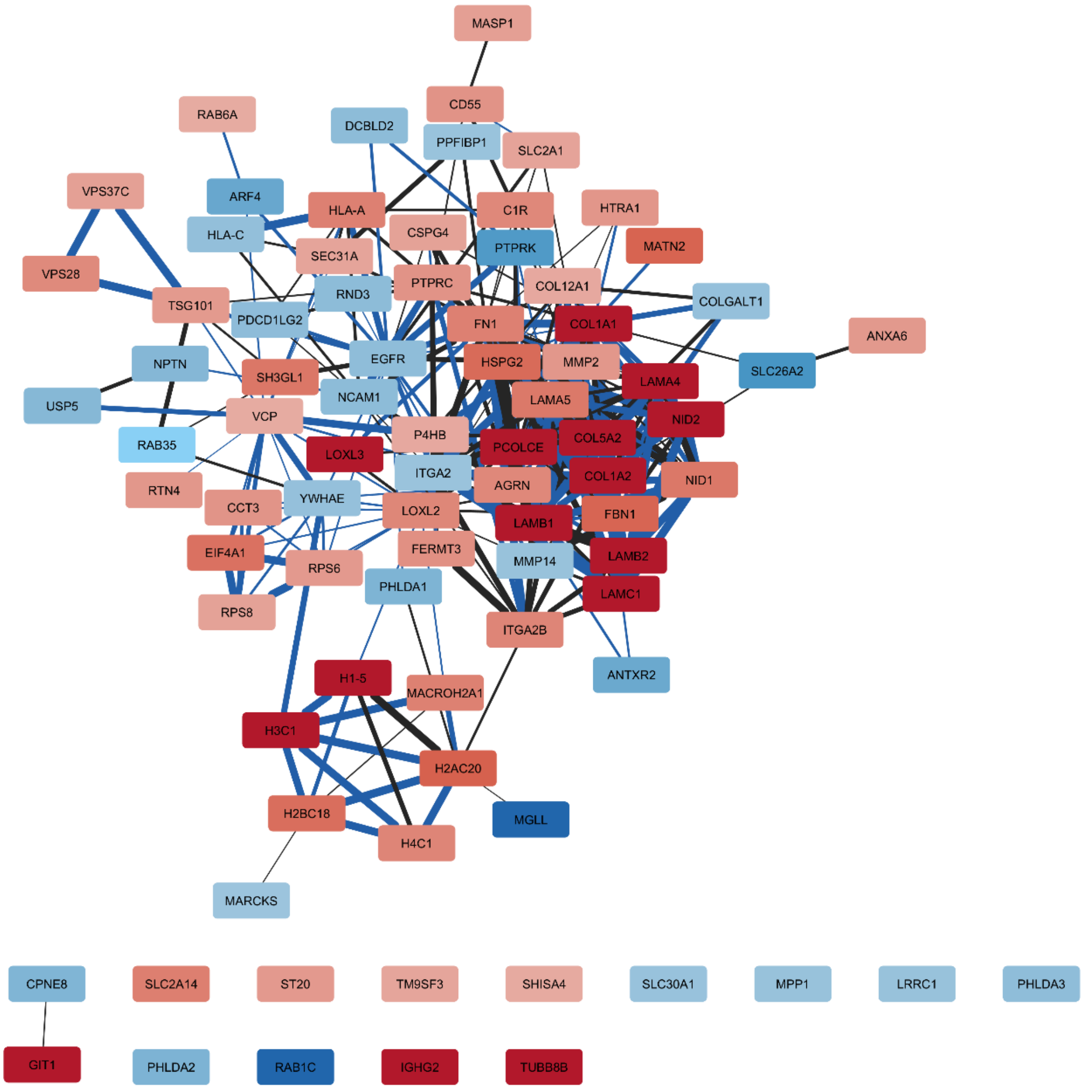
