## Supplementary Table S1 for "Effects of panitumumab on the proteome of treated EGFRvIII glioblastoma cells and on their extracellular vesicles"

**Supplementary Table S1: Gene Ontology terms and Reactome pathways enriched among differentially expressed proteins in U87-EGFRvIII-derived EVs following panitumumab treatment.**

| Category | Accession Number term | Term Name | Count | P-Value | Adj. P value (Benjamini-Hochberg) | Fold Enrichment | Gene Symbols |
|---|---|---|---|---|---|---|---|
| GO_BP | GO:0030198 | extracellular matrix organization | 15 | 0,00000239 | 0,00233 | 4,11 | MMP2,ITGA2,COL12A1,LOXL3,LAMC1,NID1,NID2,LOXL2,COL1A1,MMP14,COL1A2,COL5A2,COLGALT1,MATN2,FBN1 |
| GO_MF | GO:0005201 | extracellular matrix structural constituent | 17 | 1,51E-07 | 0,000059 | 4,30 | LAMA5,LAMB2,LAMA4,COL12A1,FN1,LAMB1,PCOLCE,LAMC1,NID1,HSPG2,NID2,COL1A1,COL1A2,COL5A2,AGRN,MATN2,FBN1 |
| GO_MF | GO:0030527 | structural constituent of chromatin | 6 | 0,0000439 | 0,00571 | 10,11 | H1-5,H2AC20,H3C1,H4C1,MACROH2A1,H2BC18 |
| GO_CC | GO:0005604 | basement membrane | 14 | 7,71E-07 | 0,0000933 | 4,80 | LAMA5,LAMB2,LAMA4,FN1,LAMB1,PCOLCE,LAMC1,NID1,HSPG2,NID2,LOXL2,AGRN,MATN2,FBN1 |
| GO_CC | GO:0031012 | extracellular matrix | 25 | 0,00000165 | 0,000119 | 2,68 | LAMA5,LAMA4,COL12A1,HTRA1,LOXL3,LAMC1,NID1,NID2,LOXL2,ANXA6,NCAM1,LAMB2,MMP2,FN1,LAMB1,PCOLCE,HSPG2,COL1A1,MMP14,COL1A2,COL5A2,CSPG4,AGRN,FBN1,MATN2 |
| GO_CC | GO:0140151 | solid phase of interstitial matrix | 11 | 0,000325 | 0,0131 | 3,65 | COL1A1,COL1A2,COL12A1,COL5A2,LOXL3,FN1,PCOLCE,HSPG2,LOXL2,MATN2,FBN1 |
| GO_CC | GO:0000786 | nucleosome | 6 | 0,000574 | 0,0208 | 7,08 | H1-5,H2AC20,H3C1,H4C1,MACROH2A1,H2BC18 |
| PATHWAY | R-HSA-1474244 | Extracellular matrix organization | 26 | 1,78E-09 | 0,00000116 | 3,45 | LAMA5,LAMA4,COL12A1,ITGA2B,HTRA1,LOXL3,LAMC1,NID1,NID2,LOXL2,NCAM1,COLGALT1,LAMB2,MMP2,ITGA2,FN1,LAMB1,PCOLCE,HSPG2,COL1A1,MMP14,COL1A2,COL5A2,P4HB,AGRN,FBN1 |
| PATHWAY | R-HSA-3000178 | ECM proteoglycans | 14 | 0,00000133 | 0,000173 | 4,52 | LAMA5,LAMB2,LAMA4,ITGA2,ITGA2B,FN1,LAMB1,LAMC1,HSPG2,COL1A1,COL1A2,COL5A2,NCAM1,AGRN |
| PATHWAY | R-HSA-1474290 | Collagen formation | 9 | 0,00024 | 0,00917 | 4,59 | COL1A1,COL1A2,COL12A1,COL5A2,LOXL3,PCOLCE,P4HB,COLGALT1,LOXL2 |
| PATHWAY | R-HSA-9818564 | Epigenetic regulation of gene expression by MLL3 and MLL4 complexes | 5 | 0,00046 | 0,0142 | 9,69 | H2AC20,H3C1,H4C1,PHLDA1,MGLL |
